# The Relationship Between Preference and Switching in Flower Foraging by Bees

**DOI:** 10.1101/2023.12.08.570807

**Authors:** Daniel R. Papaj, Avery L. Russell

## Abstract

It seems self-evident that generalist foragers switch more between resources than specialists but despite diverse ecological and evolutionary implications, how variation in switching relates to variation in preference warrants additional study. Here we tested predictions based on a simple probability model, using flower-foraging bees as a model system. In laboratory assays, we presented bumble bee (*Bombus impatiens*) workers with flowers of two species, *Tecoma stans* and *T. alata*, from which they could collect nectar and/or pollen. We quantified landing preference and occurrence of switching between species in successive visits. Bees varied greatly in floral preference. Almost half showed statistically significant preferences for one or the other species, while the rest were generalists. As expected, generalists using both flower species switched more in successive visits than bees that were more specialized, a pattern fit to a quadratic function. However, generalist individuals switched more than expected based on null expectation. A Modified Jacob’s Index (MJI) of switching was significantly positively correlated with degree of preference: generalist bees had more negative MJI’s than specialist bees, indicating that even after the expected statistical effect of preference on switching was accounted for, they switched more than specialists. A simulation ruled out the possibility that the pattern was due to bias in MJI. Generalist-specialist differences in which food was collected (nectar versus pollen) were also ruled out. We discuss possible explanations for our observed pattern and outline possible ways in which preference and switching frequency will interact to shape pollinator behavior and the floral resource.

**Significance statement:** Behavioral preference is the subject of a large literature in areas such as foraging, mating and communication. However, a preference measure alone does not necessarily tell us if choices for one alternative are made in runs or intermingled with choices for another alternative. The distinction between preference and the sequential pattern of choices is relevant in many contexts in behavioral ecology but has been a particular focus of study in flower foraging by pollinators. Even in that literature, the relationship between preference and sequential pattern in switching warrants further examination. In our study, bees were shown to vary in preference for flowers of two species. Some were generalists; some were specialists on one or the other species. Generalist bees switched more than specialist bees, even after controlling for statistical effects of preference on switching frequency. The report of this generalist-specialist pattern in switching may be novel and has far-reaching implications throughout the field of behavioral ecology.

## Introduction

The measurement and analysis of resource preference is one of the pillars of the field of behavioral ecology (Emlen 1966; Manly 1972; Chesson 1978; Krebs 1973; Stephens and Krebs 1985; Manly et al. 2007; Abbott and Dukas 2016; Michelot et al. 2019). Likewise, the analysis of resource switching by foragers has long received attention in theory and experiments (Bateman 1951; Murdoch 1969; Abrams 2006; Hooker et al. 2017; Fathipour et al. 2020). We might intuitively expect the frequency of switching between resource types to relate to some extent to a forager’s pattern of preference for the resource. A forager that strongly prefers one resource type over others will tend to make more successive visits to that type by chance than a forager that shows relatively weak preference. Such linkage is important to consider because preference and switching may be subject to different selection pressures. For example, one could imagine that a forager’s preference might be influenced most by nutritional requirements (Raubenheimer and Simpson 2018), while its propensity to switch between even two identical resources might be affected most by constraints on learning and memory (Chittka et al. 1999). If preference and switching are intrinsically linked, selection for dietary mixing might affect switching, and learning and memory constraints might affect preference.

Resource preference and switching have been studied in the context of plant-pollinator interactions for additional reasons. Pollinator behavior, including innate and learned cue preferences, play a key role in driving the evolution of floral traits (Schiestl and Johnson 2013; Haverkamp et al. 2018; Finnell and Koski 2021; Lanterman et al. 2023). Pollination biologists have also considered how pollinators switch among plant species as they forage because switching has implications for the effectiveness of pollen transfer (Bateman 1951; Waser 1983, 1986; Janovsky et al. 2017; Bruninga-Socolar et al. 2023).

In this study, we sought to assess how switching varies with preference, using flower foraging by bumble bees as a model system. We begin this assessment by proposing a simple probability model of how preference and switching might be intrinsically related. In this model, the probability of switching from a plant type A to a plant type B at each step in a foraging bout depends strictly on the overall preference for the two plant types. The predicted pattern of switching is the quadratic function of the forager’s overall preference shown in Fig. 1a. Non-preference is associated with the greatest probability of switching, with values declining in non-linear fashion as preference for one or the other plant type becomes increasingly stronger.

**Fig. 1.**
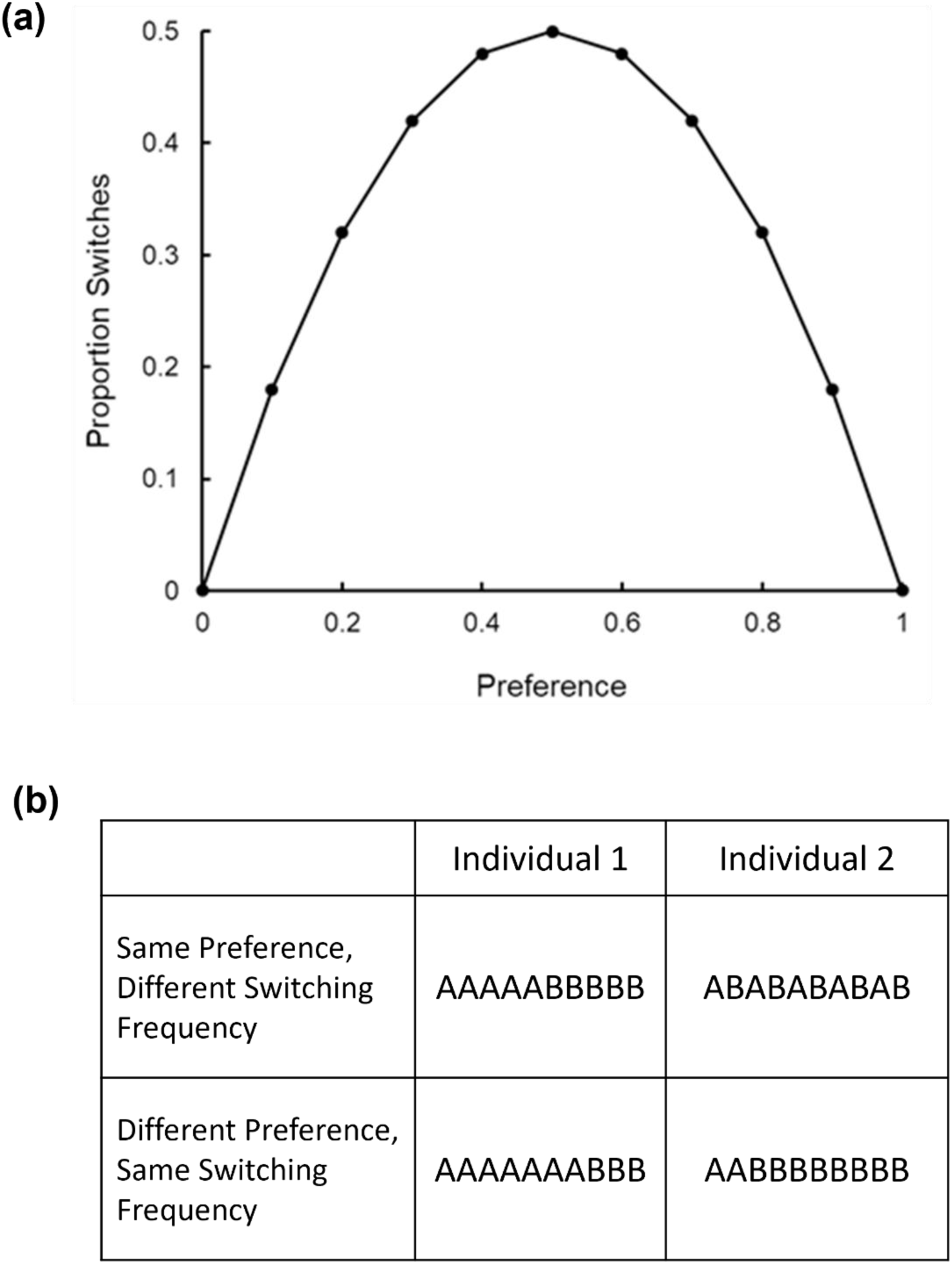
(a) Null expectation for the proportion of switches between two resources A and B over preference, which fits to a quadratic function. Proportion switches was calculated as [1 – (P_A_^2^ + P_B_^2^)] where P_A_ is the probability of using resource A and P_B_ the probability of using resource B. P_A_^2^ is the probability of an A-A transition and P_B_^2^ the probability of a B-B transition. (b) Across the first row are two hypothetical individuals having the same overall preference but different switching frequency. Across the second row are two hypothetical individuals differing in preference but having the same switching frequency.

We consider this pattern to be a null expectation for the relationship between preference and switching. However, foragers may show a different pattern of switching and preference. Foragers can conceivably have the same preference, or even lack of preference, yet differ in their degree of switching (Fig. 1b). Consider two individual hypothetical pollinators, neither of which show an overall preference for one of two plant species A and B. The two plant types are intermingled in the pollinators’ search paths, and 50% of their visits are made to each plant type. The first individual shows fidelity first to one type and then to the other in its visits: AAAABBBB. The second individual switches systematically, alternating visits between the two plant types: ABABABAB (Fig. 1b, first row). Even though the two individuals do not differ in overall preference, the first individual is potentially a better pollinator of each plant species, because it more often visits the same species successively and is thus more likely to transfer conspecific pollen and less likely to transfer heterospecific pollen.

Individuals can likewise differ in preference yet show similar patterns of switching (Fig. 1b, second row). Consider an individual pollinator that switches minimally, and strongly prefers A in a series of visits: AAAAAAABBB. Compare that to an individual that also switches minimally but strongly prefers B: AABBBBBBBB. Both pollinators show the same frequency of switching but preference is different in sign and magnitude. These hypothetical examples illustrate that switching frequency may in principle deviate from null expectation in ways that have ecological relevance for plant and pollinator.

Various indices have been developed to assess the extent to which switching frequency deviates from null expectation based on preference (Bateman 1951; Jacobs 1974; Waser 1986; Gegear and Thomson 2004), and to explore causal factors for deviations found. However, the indices have not, to our knowledge, been used to assess precisely how the propensity to switch is related to individual variation in the overall pattern of preference. Using such an index, we examined how the tendency of individual bumble bees (*Bombus impatiens*) to switch between flowers of two plant species was related to their preference for one or the other species. We predicted that switching frequency would vary with preference in the form of the quadratic function shown in Fig. 1a. Because the two plant species used are closely related, we predicted most bees would be generalists, showing modest preference at best. However, despite highly controlled laboratory assay conditions, we found that initially naïve bees varied substantially in floral preference. Some bees specialized on one species, some on the other, and some were generalists. The variation in preference allowed us to examine how switching frequency varied with degree of specialization.

## Methods

### Plants

We studied preference and switching by bees with respect to two species of *Tecoma* (family Bignoniaceae), the flowers of which are locally available in the Tucson area over much of the year. The first was a variety of *Tecoma stans* (L.) Juss. ex Kunth. *Tecoma stans* is commonly known as “yellow trumpetbush” or “yellow bells,” and like other species in the family, produces sympetalous flowers with petals fused to form a tube-like corolla. The species is globally distributed and native to parts of the southern United States, including Arizona and Florida. Over its range, it is visited by bees, butterflies, moths, and hummingbirds (Pelton 1964), but is generally characterized as bee pollinated. The species has been cultivated as an ornamental; flowers of *T. stans* plants growing on the University of Arizona campus resemble a variety called Gold Star. Flowers on campus have been observed to be robbed by carpenter bees (*Xylocopa sonorina* and *Xylocopa californica*), and also secondarily robbed and legitimately visited by honey bees (DRP and ALR, unpubl. obs.). Flowers of the native *T. stans* occurring in the nearby Santa Rita Mountains south of Tucson are visited by carpenter bees (*X. californica* and *Xylocopa micheneri*) as well as by the Sonoran Bumble Bee, *Bombus sonorus*. It is also visited by solitary bees (for example, *Anthophora* spp.) (DRP and ALR, unpubl. obs.). Hummingbirds also visit *T. stans* (Pelton 1964; DRP, unpubl. obs.). The *T. stans* flowers used in the experiment were taken from plants on the University of Arizona grounds.

The taxonomic status of the other *Tecoma* used in our study is uncertain. It is sometimes listed in the nursery trade as a pure species, *Tecoma alata*, a synonym of *Tecoma guarume*. It may also be referred to as ‘Orange Jubilee’ and described as a hybrid between *T. alata* and *T. stans.* There are yet other attributions in the horticultural literature (Meerow and Ayala-Silva 2008). We will refer to it here as *T. alata*, its most commonly used name. In urban settings in southern Arizona, where it is a common ornamental planting, *T. alata* is visited by hummingbirds. Although bumble bees are rarely observed in urban Tucson, honey bees and two carpenter bee species (*X. californica* and *X. sonorina*) are frequent visitors.

In our assays, we used flowers from inflorescences that had been covered with organza bags for at least one full day on plants of either species on the University of Arizona campus, to prevent visitation by insects and birds. Flowers were used the same day they were picked.

### Pollinator

As a model pollinator, we used the commercially available Eastern Bumble Bee, *Bombus impatiens*, which is common throughout eastern North America, from Maine to Florida and west through Ohio (Discover Life). In its native range, the species is a generalist visiting a wide array of flowering plant species offering nectar and pollen, including *Tecoma stans*. *Bombus impatiens* has been used extensively in studies of bumble bee behavior.

Colonies were purchased from Koppert Biological Systems (Howell, MI). Approximately equal numbers of workers from 3 colonies were used in the experiment. Colonies foraged for pollen and nectar within an 82cm × 60cm × 60cm (L x W x H) feeding chamber made from plywood and painted neutral gray on the interior. The chamber had clear acrylic ceilings and was lit from above 40W 4400 lumen LED lights (2 x 2 LED Panel; 5000K Cool White, James Industry).

For maintenance, sucrose was provided as a 20% w:w solution dispensed to bees via braided cotton wicks (6 inch Braided Cotton Rolls, Richmond Dental) extending into 40 dram vials (BioQuip Products, Inc.). Pulverized honey bee-collected pollen (Koppert Biological Systems) was dispensed on chenille fibers glued to the inside of 40-dram vials (Russell and Papaj 2016).

### Testing

Testing took place in an arena identical in design to the feeding chambers. Flower-naïve bees were captured in vials at sucrose dispensers in the chamber and then released in the test arena. This protocol ensured that bees were foragers, and in a physiological reward collection state at time of testing. The testing arena contained a vertical array of 8 previously unvisited flowers, four of *T. stans* and four of *T. alata* (Fig. 2). Flowers were mounted with pedicels inserted in custom-built water tubes that were Velcro-mounted to a 60cm x 60cm wall at one end of the arena. Flowers were spaced 7cm apart at points along the perimeter of a square. Flowers of the different types were systematically alternated by position (Fig. 2). Two such arrangements are possible wherein one or the other species were positioned on the corners of the grid; both arrangements were used the same number of times and alternated systematically in time across bee trials. No effect of arrangement on preference was detected and this factor was not included further in statistical analysis.

**Fig. 2.**
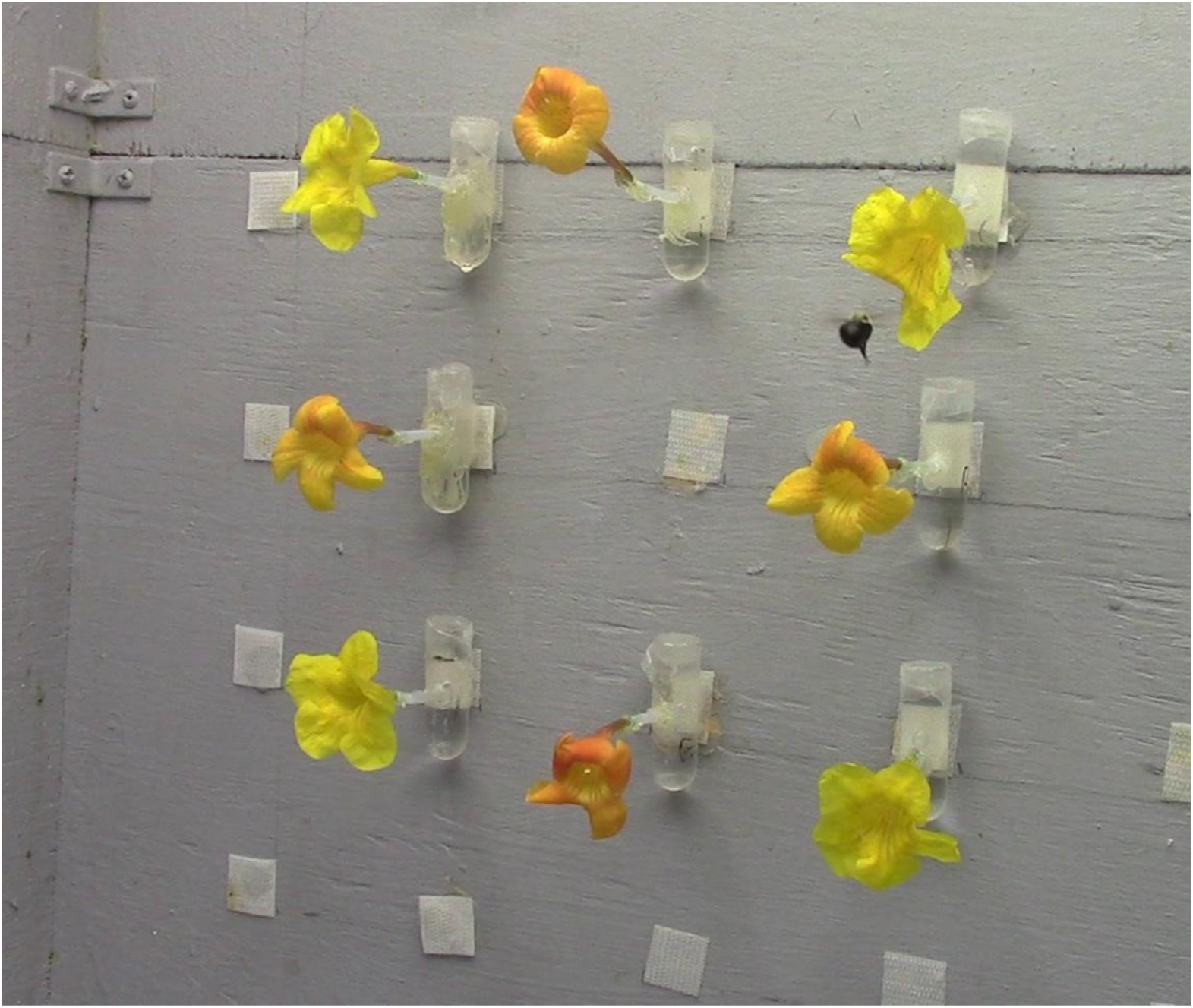
Photo of vertical array showing 4 *Tecoma stans* (yellow flowers), 4 *T. alata* (orange flowers), and 1 test bee in the test arena. Flowers are mounted in customized water tubes that are Velcro mounted to the back wall of the arena. Along the square formed by the flowers, flowers are spaced 7 cm apart on the horizontal and vertical axes.

Flower-naïve bees were released individually into the foraging arena and allowed to visit the *Tecoma* flower array. A visit was defined as the bee landing on a flower (contact of at least 4 legs with the flower). After 10 minutes elapsed without a visit, we terminated the trial and euthanized the bee to prevent it from returning to the colony and influencing the foraging behavior of other bees. The number of visits in a single trial ranged from 9 to 75, but most bees made ca. 40 visits (mean ± SE number of visits: 35.7 ± 3.1; N=23). Flowers were discarded after the trial and replaced with fresh, initially unvisited flowers.

We recorded if bees attempted to collect a floral reward during a visit. Bees can obtain nectar and/or pollen from either *Tecoma* species. These food types are similarly segregated spatially within each species’ flowers, which allowed us to distinguish which were being collected during a visit. A bee was recorded as attempting to collect nectar if it crawled deep into the corolla past the anthers, where the nectary was located. Prior to this study, we confirmed that bees searched for nectar by shining a light on the flowers for a subset of bees and observing that the bees’ probosces were invariably extended in silhouette (DRP, unpubl. obs.). A bee was recorded as attempting to collect pollen if the bee sonicated the anthers. Sonication is a behavior in which pollen-foraging bees use substrate-borne vibrations generated by contraction of thoracic muscles to dislodge pollen from anthers (Buchmann 1985). *Bombus impatiens* reliably sonicates when collecting pollen from *Tecoma* flowers. The sound accompanying the vibrations is readily perceptible to human ears. We noted occurrences of sonication at flowers and inspected the bees’ pollen baskets (corbiculae) for presence of pollen.

Each visit to a flower was classified as one of the following four types: pollen collection attempt only, nectar collection attempt only, both pollen and nectar collection attempts, or neither nectar nor pollen collection attempts.

To minimize observer bias, behavioral data collection was done blind. Observers did not know the primary aims of the experiment or how the data would be analyzed.

### Analysis of switching

We used two approaches to analyze the frequency of switching, estimated as the proportion of pairwise transitions that were heterospecific. First, a quadratic function was fit to switching frequency over the range of individual preference values (see Fig. 1a for the quadratic function expected under a null model of switching).

Second, we used a modification by Gegear and Thompson (2004) of an index developed by Jacobs (1974):

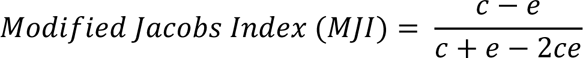

where ‘c’ is the proportion of moves between the same species, and ‘e’ is the expected proportion of moves between the same species based on the overall frequency of each species visited in a trial. The Modified Jacobs index (MJI) controls for bias in the frequency of visits to each species. MJI can range from −1 (switching as much as possible given preference) to 0 (random pattern of switching in relation to preference) to +1 (switching as little as possible given preference). Two of our 23 test bees showed an absolute preference for *T. stans*; for those bees, MJI is undefined and, to be conservative in our analysis, these bees were excluded from analyses of MJI and preference.

### Simulation of preference and switching

To confirm that MJI is not biased by overall preference, we conducted a stochastic simulation. Using Excel (Microsoft, Inc., version 365 ProPlus), we designated 5 bins of mean preference (as a proportion) for one of two hypothetical plant species (A and B), ranging in steps of 0.1, from 0.5 (non-preference) to 0.9 (strong preference for A or B). Each of 10 bees simulated within each bin made 100 visits to Species A or Species B. The species visited at each step was determined by the *rand* function in Excel, with the probability of a visit being made to Species A or B depending on the preference bin to which the bee belonged. Thus, a bee assigned to the 0.7 bin had a 0.7 chance of visiting Species A and a 0.3 chance of visiting Species B. We then computed the overall preference and MJI for that bee from its string of 100 visits and repeated this for 9 more bees within a preference bin. The protocol generates stochastic variation among bees in each preference bin both for preference and for MJI, while constraining preference to vary from non-preference to strong preference within the simulated population.

### Analysis of collection of floral rewards

To determine if preference and/or switching were related to the type of nutrient (nectar and/or pollen) that individuals were collecting, we quantified the profile of food collection attempts of each bee on each species of flower visited, using the following equation:

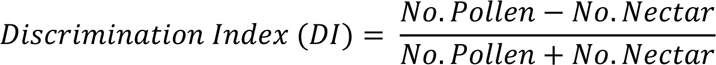

No. Pollen refers to the total number of flower visits in a trial that included an attempt to collect pollen and No. Nectar to the total number of flower visits that included an attempt to collect nectar. A DI value of 1 indicates pollen collection only, a value of −1 indicates nectar collection only, and a 0 indicates collection of both foods at equal frequency. We compared DI’s on *T. stans* with those on *T. alata* for those bees that visited both species.

We endeavored to confirm that food was being collected when attempts were made. The accumulation of pollen in the corbiculae (pollen baskets) was apparent for every bee that sonicated test flowers, confirming that pollen was collected when flowers were sonicated. Nectar collection is more difficult to confirm. Nectar is stored in the crop, and this organ may already be partly filled when the bee begins foraging. To confirm that bees were collecting nectar and to characterize how much nectar was collected, we compared nectar volumes in flowers presented to bees in assays with control flowers from the same inflorescences. Control flowers were collected at the same time as test flowers and held under the same conditions in the lab. We used microcapillary tubes (Drummond Scientific, Inc., Broomall, PA) to collect nectar and estimate volume. Measurements of both test and control flowers were made immediately following the assays in which test flowers were used. We compared control flowers to test flowers for a total of 12 test bees. Finally, we measured nectar concentration of a sample of flowers of each species using a Brix refractometer (Bellingham & Stanley Ltd., Kent, U.K.).

### Statistical analysis

All statistical analyses were made using JMP software (version 14; SAS Inc.). We used nonparametric tests such as Wilcoxon signed rank tests to analyze overall species preference and chi-square goodness of fit tests to analyze individual species preference. We used a general linear model to assess the relationship between preference and relative frequency of switching. Because bouts varied in total number of visits which might conceivably affect preference and switching, the total number of visits in the trial was included as a factor in the analysis. We used linear regression to evaluate the relationship between MJI and degree of preference. In exploratory analyses, we included colony ID as a factor. However, the colony effect was never significant, and this factor was not included in the final models. Finally, we used a repeated measures ANOVA to assess whether individual bees were collecting different foods from the two species.

## Results

### Species preference

Most bees (21 of 23) visited both species at least once. Individual species preference ranged from absolute *T. stans* preference to nearly absolute *T. alata* preference (Fig. 3; proportion visits to *T. alata*, range = 0 −0.96). Overall, bees showed a species preference for *T. stans,* making approximately 60% of their visits on average to that species (mean proportion visits to *T. alata* = 0.398 ± 0.006 S.E., N = 23). However, this mean preference was not statistically different from 0.5 (Wilcoxon signed-rank test, W = −57.5, p<0.08), reflective of substantial variation among bees.

**Fig. 3.**
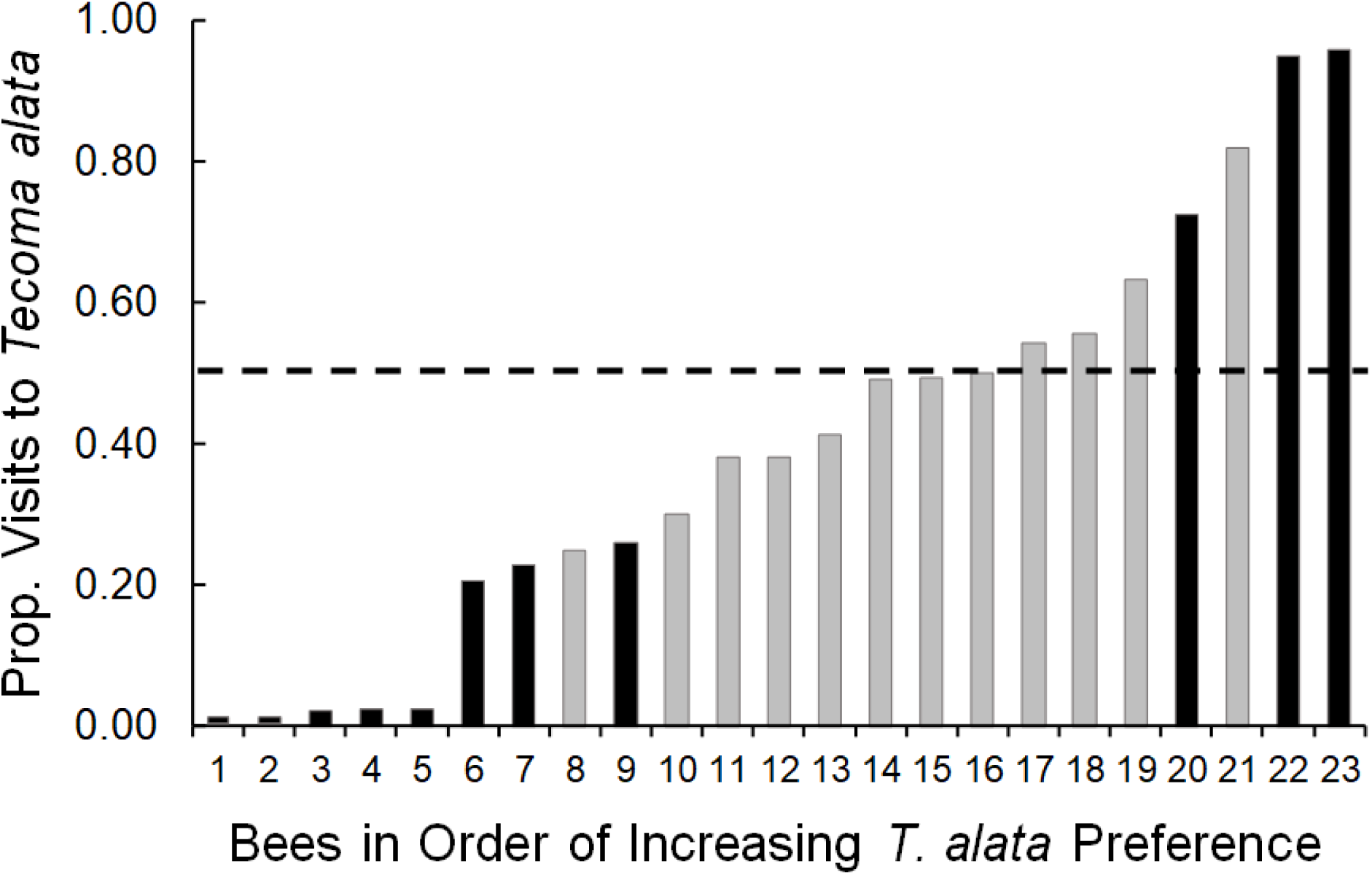
Distribution of species preference by initially naïve *Bombus impatiens* workers, expressed as the proportion of visits in the test bout made to *Tecoma alata*. Dashed line at prop. visits = 0.5 indicates non-preference. Black bars indicate values that were significantly different from 0.5 at p < 0.002 or less, according to a chi-square goodness-of-fit test. Gray bars indicate values that were not significantly different. See Online Resource 1, Table S1 for frequency counts and chi-square statistics for each bee.

Eleven of 22 bees that made enough visits for analysis showed a preference that was statistically significantly different from 50% (Fig. 3; Online Resource 1, Table S1). Eight of those 11 bees preferred *T. stans* to a significant extent while 3 bees significantly preferred *T. alata*. The remaining 11 bees visited both species but did not show a statistically significant preference. Of those bees, four were absolute generalists, making a virtually identical number of visits to each species (Fig. 3; Online Resource 1, Table S1).

### Switching frequency and preference

Consistent with null expectation (Fig. 1a), the proportion of switches in pairwise transitions was greater at intermediate values of preference (Fig. 4) than at more extreme values. A quadratic fit to the data (solid line) explained a large and significant proportion of the variance (Fig. 4; R^2^ = 0.93; F_2,20_ = 123.87, p<0.0001).

**Fig. 4.**
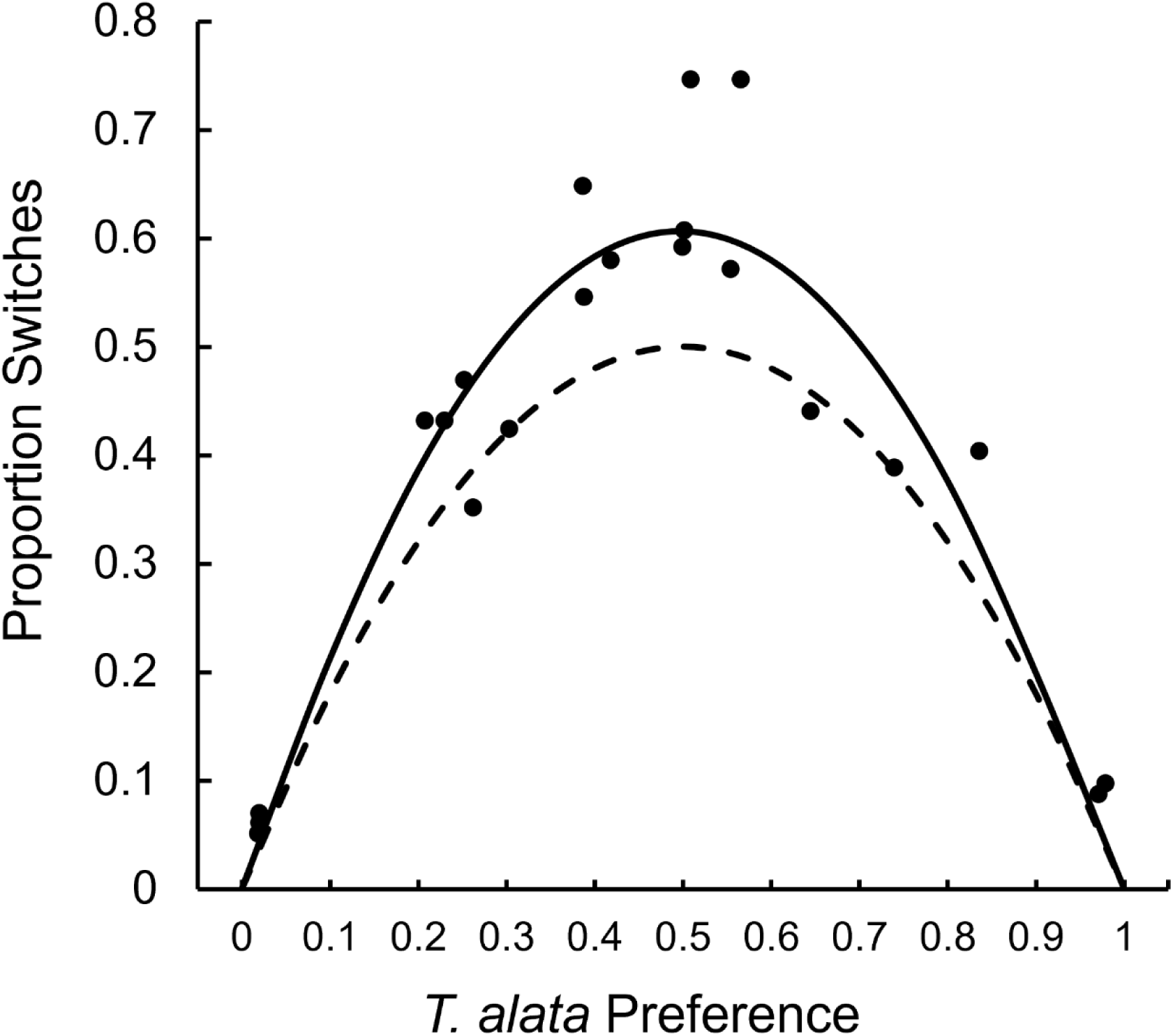
Quadratic fit to the proportion of pairwise transitions that involved a switch between plant species, in relation to species preference, estimated as proportion of visits made to *Tecoma alata* (solid line). For comparison, the function based on the simplest null expectation is also shown (dashed line, based on probability model, see Introduction and Fig. 1a). Solid bullets indicate individual data, with some lying on top of each other (Total N = 22; data presented in Online Resource 1, Table S1).

Despite the strong fit to a quadratic model, data did not precisely meet null expectation. When the parabola based on null expectation is superimposed on the parabola fit to the data (Fig. 4), bees appear to be switching more frequently than expected, as indicated by the space between the two parabolas. This difference is greatest at values of non-preference, i.e., for generalist bees, and then decreases as preference becomes closer to 0 (specialization on *T. stans*) or closer to 1 (specialization on *T. alata*). This result implies that even after we account for the effect of preference on the frequency of switching, generalists are switching more than specialists.

To determine if these inferences were statistically robust, we computed a Modified Jacobs Index (MJI) for each test bee (see Methods). Values for the Modified Jacobs Index (MJI) ranged from −0.543 to 0.077, positive values reflecting less switching than expected in relation to preference and negative values reflecting more switching than expected in relation to preference. Mean MJI overall was significantly less than zero (mean MJI = −0.157 ± 0.039 S.E.; Wilcoxon signed-rank test against expected mean of 0, W = −91.5, p<0.0003; N=21), indicating that bees overall alternated visits to the two species more than expected based solely on their preference.

We next assessed how the index changed with degree of preference. Degree of preference was formulated as the absolute value of the deviation of proportion visits to *T. alata* from 0.5. A value of 0 for this metric indicates a pure generalist; a value of 0.5 indicates a pure specialist for one or the other species. A general linear model was used to assess how MJI changed with degree of preference as well as with the total number of visits in a trial. Our model explained a significant amount of the total variance (R^2^=0.58; F_2,18_=12.32, p=0.0004). The model intercept was significantly less than 0 (estimate of −0.52; t_1,18_=6.58, p<0.0001). MJI became significantly less negative with increasing degree of preference (F_1,18_=12.44, p<0.003; β=0.0057 ± 0.0016 S.E.; Fig. 5a) and significantly less negative as total number of visits increased (F_1,18_=14.15, p<0.002; β=0.0066 ± 0.00176 S.E.; Fig. 5b). Thus, generalist bees switched disproportionately more than specialist bees, even after we accounted for the expected effect of preference on switching.

**Fig. 5.**
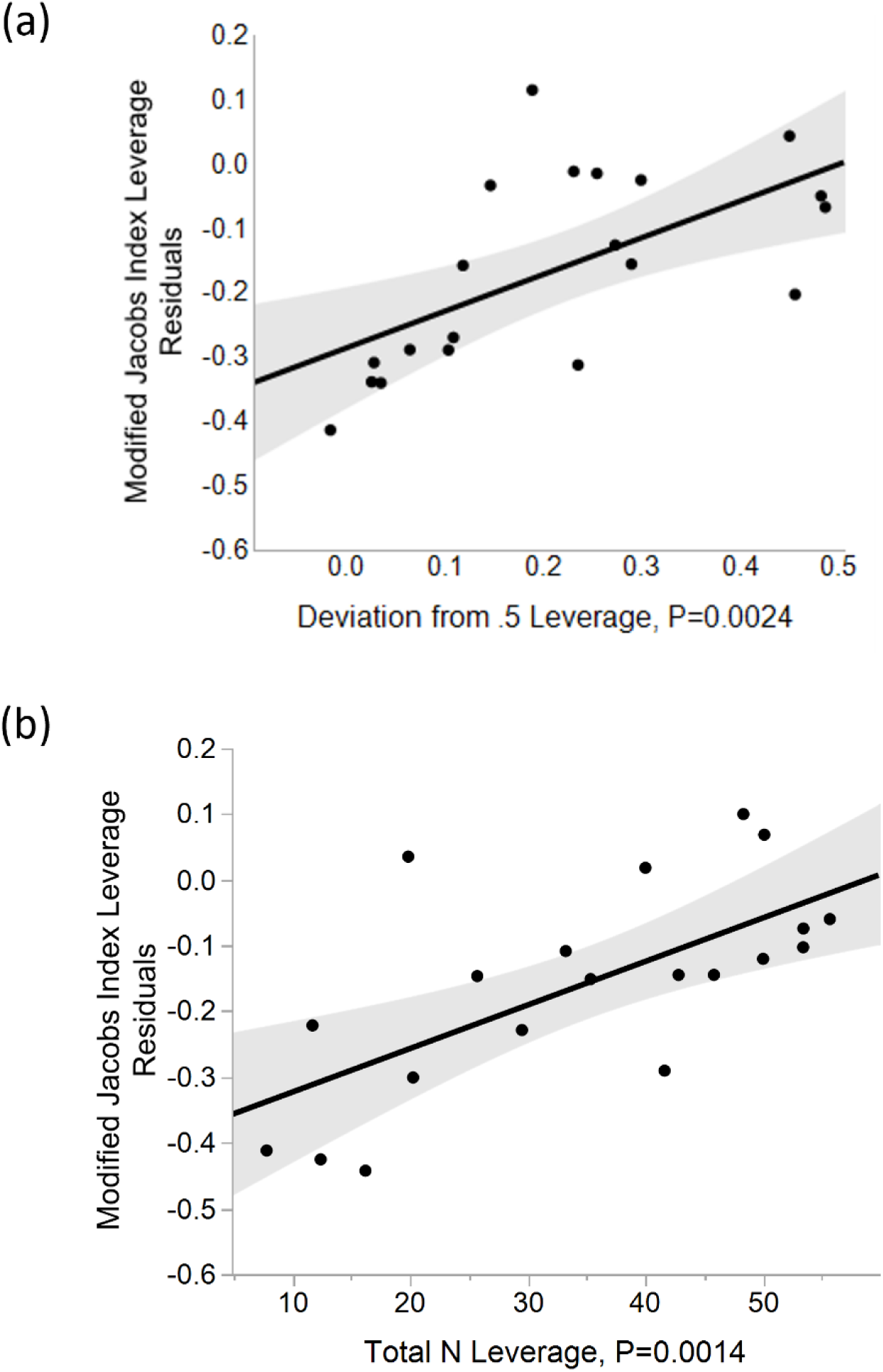
Leverage plots of residuals for a general linear model with Modified Jacobs Index (MJI) as the dependent variable. (a) A measure of preference (deviation from a preference value of 0.5) as independent variable. Larger values of deviation indicate stronger preference. (b) Total number of visits in a bout (N) as independent variable. Line in each panel indicates model fit. The shaded region indicates 95% confidence band. Solid bullets show residual values based on fit to the model.

### Simulation of Modified Jacobs Index in relation to preference

One possible explanation for the disproportionately greater switching of generalist bees, relative to specialist bees (Fig. 5a), is that it is an artifact of the index used. To evaluate the possibility that MJI is arbitrarily biased and intrinsically positively correlated with preference, we conducted a stochastic simulation. The simulation assumes that foragers switch relative to preference according to the null model presented in Fig. 1a (see Methods). Results of the simulation are shown in Fig. 6. Simulated MJI values are distributed symmetrically around a value of 0 over the range of preference simulated, confirming that MJI is not intrinsically positively correlated with preference. In short, over the range of preference values assessed, MJI appears to be an unbiased estimator of degree of switching in relation to preference.

**Fig. 6.**
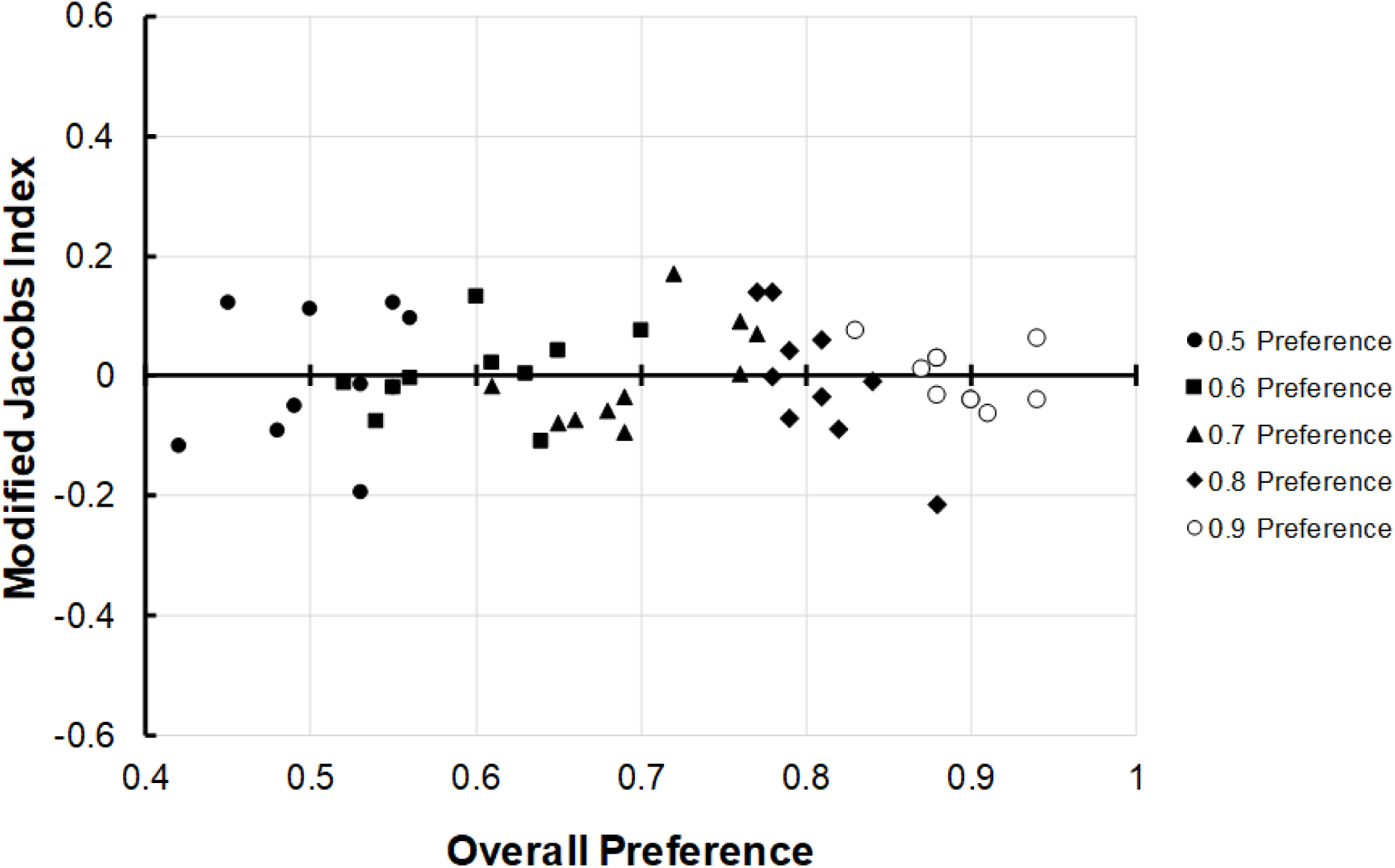
Results of a simulation of the Modified Jacobs Index (MJI) in which both overall preference and occurrences of switching vary stochastically within each of 5 preference bins. Bullet symbols indicate the assigned preference bin for each value of preference. Different bullets in the same bin illustrate stochastic variation in simulated MJI within that bin.

### Patterns of food collection

Another possible explanation for the disproportionately greater switching of generalist bees relative to specialist bees, is that generalist bees are actively mixing their diet in time, and alternating collection of one food (nectar or pollen) from one *Tecoma* species with collection of the other food from the other species. No such pattern was observed.

Fourteen bees visited both *Tecoma* species. The relative use of pollen and nectar on the two species was assessed with a Discrimination Index (DI; see Methods). A DI value of 1 indicates pollen collection only, a value of −1 nectar collection only, and a 0 indicates collection of both foods at equal frequency. The possibility that bees collect different foods from the two *Tecoma* species depending on their degree of species preference was evaluated with a repeated measures ANOVA analysis of Discrimination Index (DI) on *T. stans* versus DI on *T. alata*, including degree of preference as a factor. A significant effect of preference in the model would indicate that the difference in DIs for each bee depended on preference. In analysis of 14 bees collecting foods on both species, mean DI on each *Tecoma* species was slightly negative, indicating a slight bias towards nectar collection on both species. Mean DI values were very similar on the two species (−0.116 on *T. alata* and −0.144 on *T. stans*). There was no significant interaction between the difference in DI on the two *Tecoma* species and degree of preference (F_1,12_ = 0.007, p=0.93). In short, generalists appear to be collecting foods in roughly the same way on the two species as more specialized bees are. Moreover, the relative frequency of pollen collection attempts on *T. stans* was positively correlated with the relative frequency of pollen collection attempts on *T. alata* (Fig. 7; Spearman’s Rank Correlation, ρ = 0.769, p=0.0013).

**Fig. 7.**
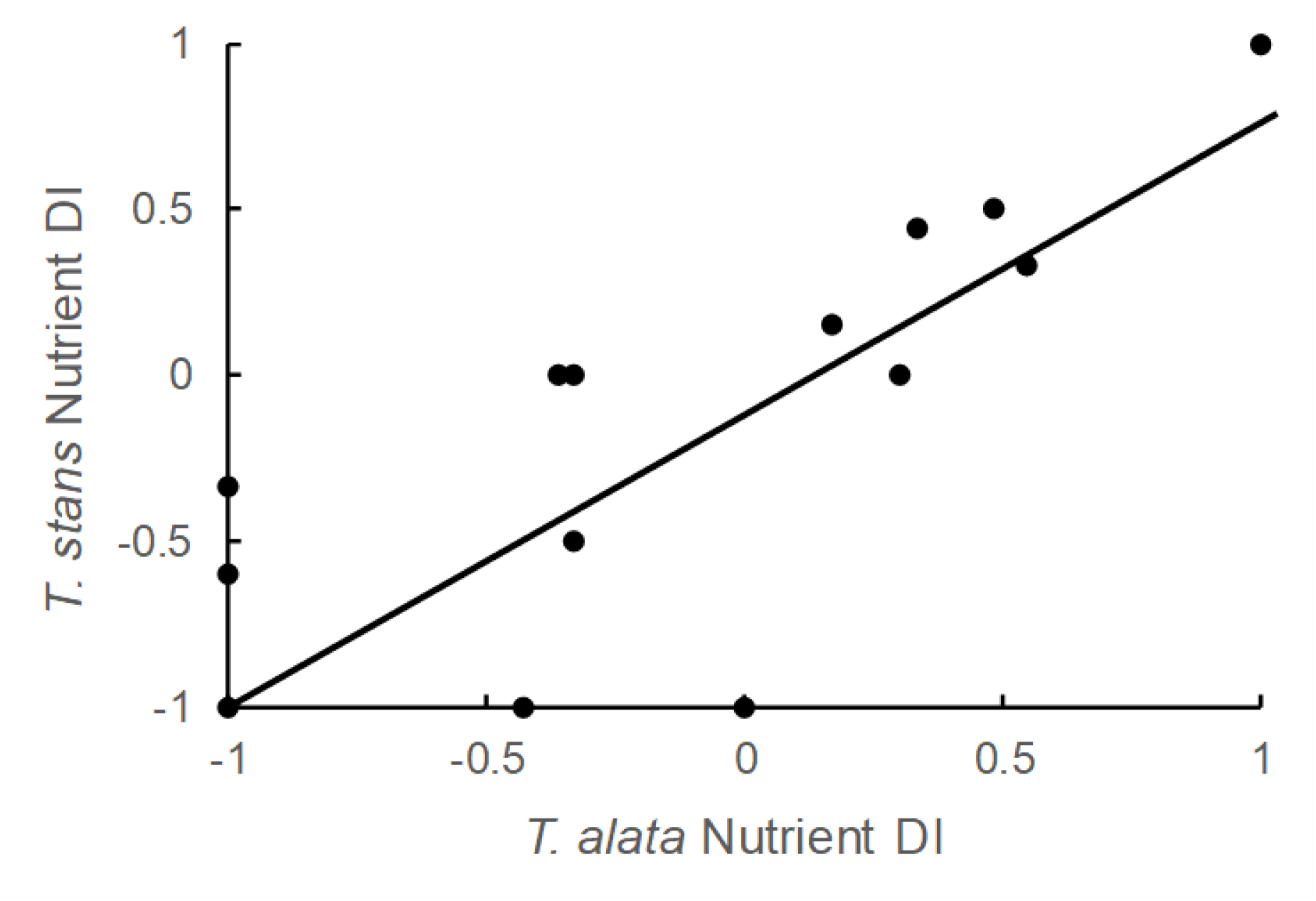
Relationship between foods sought (pollen and/or nectar) on *Tecoma stans* and foods sought on *Tecoma alata* for test bees that visited both species. DI stands for Discrimination Index (see Methods for details). A value of 1 for a DI on a given species indicates that only pollen was sought on that species; a value of −1 indicates that only nectar was sought on that species. A value of 0 along either axis means a bee made an equal number of attempts to collect pollen and nectar in its trial. Solid line indicates best fit in a linear model. N (bees) = 14.

### Analysis of nectar collection

Analysis of nectar collection confirmed that bees making nectar collection attempts were successfully extracting nectar from both *Tecoma* species (Online Resource 1, Table S2; see Methods).

## Discussion

### An intrinsic correlation between preference and switching frequency

Our study found that floral preference and switching frequency were correlated roughly as expected under a simple probability model that presumes these traits to be intrinsically related (Fig. 1a). Generalists switched more than specialists, the pattern fitting a quadratic function. To the extent that preference and switching are intrinsically related, selection on one trait will necessarily influence evolution of the other trait. Selection for high switching frequency, in the absence of selection on preference, should result in the expression of weak preference. Selection for strong preference, in the absence of selection on switching frequency, should result in expression of low switching frequency.

While direct evidence of patterns of selection specific to one or the other trait is scant, such specificity is implicit in some discussions of preference and switching. For example, some treatments of so-called floral constancy in pollinators propose that near-absolute floral preference results from strong selection for low switching frequency. The learning investment hypothesis (Laverty 1994; Grüter and Ratnieks 2011) proposes that switching incurs costs associated with learning or re-learning routines for removing floral rewards. Strong preference results from selection against switching. A competing costly information hypothesis (Chittka et al. 1999; Grüter and Ratnieks 2011) proposes that acquiring information about rewards of novel floral species is costly and should be avoided if the current floral resource is sufficiently profitable. In this case, strong preference will result due to sampling costs associated with switching to a novel species.

The reverse may be true, such that switching frequency is affected by selection on preference. For illustration, Bernays and colleagues proposed that herbivores evolve specialization for host plants because a close association with a given host species facilitates the evolution of defense against natural enemies (Bernays 1988, 1989; Bernays and Graham 1988). If selection against predation favors host-specific crypsis, for example, selection will favor specialization on a host while not acting directly on switching frequency. Yet due to the intrinsic correlation between preference and switching, the specialist would be expected to show relatively little tendency to switch between host species.

### Tradeoffs between preference and switching

The scenarios above consider selection on one trait as though the other trait was not itself under selection. It is possible, perhaps likely, that selection acts on each trait simultaneously. Selection could be congruent or opposing (Table 1). Where selection is congruent, selection on preference will reinforce selection on switching frequency, and vice-versa. The effects of congruency of selection may be the most difficult to document because preference and switching are expected by default to be related independent of selection.

**Table 1.**
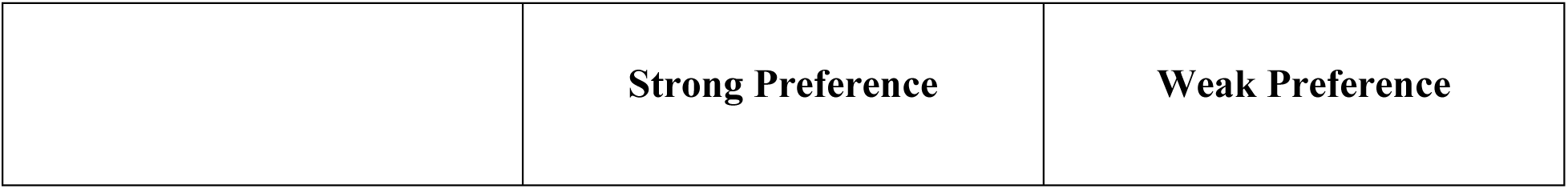
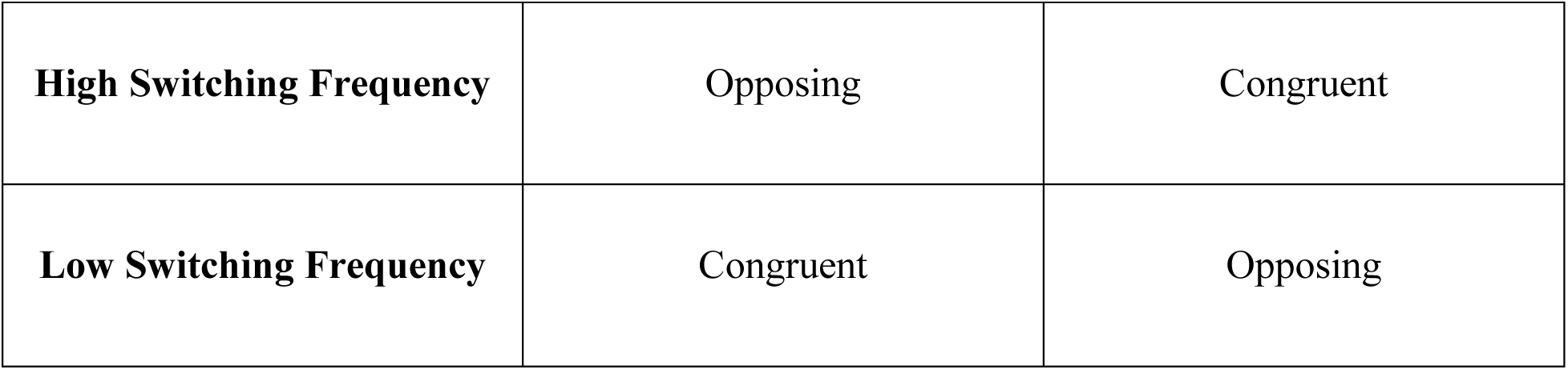
Possible patterns of congruency of selection on preference and switching frequency.

Selection on preference and switching frequency can also act in opposition. Selection for strong preference and selection for high switching frequency oppose each other, as does selection for weak preference and selection for low switching frequency (Table 1). In these circumstances, there can be two outcomes. First, the tradeoff may favor one or the other pattern, depending on associated fitness consequences. For example, Hooker et al. (2017) reported a tradeoff between food preference and switching. They found that Arctic charr performed better on a mixed diet than on a single diet item yet specialized on a single item, a result they attributed to high costs of switching.

Alternatively, selection may act to weaken the correlation between preference and switching frequency, resulting in some of the patterns mentioned in the Introduction (Fig. 1b) where preference and switching frequency are mismatched. A widely accepted characterization of floral constancy implies such a mismatch, wherein pollinators “sometimes restrict their visits to flowers of a single species or morph within a species, *even when rewarding alternative flowers must be bypassed in the process*” (quoted from Waser 1986; italics added). The implication of such a pattern is that a pollinator might do better to relax its preference and reduce travel costs, but does not, presumably because selection against switching disfavors it (Chittka et al. 1999).

### Interaction between preference and switching

Possibly the most intriguing finding in our study was the generalist-specialist pattern in switching. By default, we expect generalists to switch more than specialists (Fig. 1a). However, generalist bees in our study were observed to switch more, even after adjusting for the expected effects of preference on switching (Fig. 7). To our knowledge, the graded relationship we observed has not been reported previously in the foraging literature.

We make no claim at this point as to how general the relationship between switching frequency and degree of preference is. It is possible, for example, that the relationship is sensitive to parameters of the assay. We used an array in which flowers of the two species were alternated systematically along the perimeter of a square. It is conceivable that generalist bees were ‘traplining’ (Thomson et al. 1987, 1997; Klein et al. 2017) through the array differently than were specialist bees, with a greater tendency to encounter flowers of different species within their search path. Many studies of floral choice under controlled conditions alternate the positions of flowers in space, to control for location effects in such assays (Ohashi and Thomson 2009; Reynolds et al. 2013). It would be interesting to manipulate the spatial distribution of the two species in an array to determine if and how distribution affects the preference-switching relationship.

One factor we considered as possibly important in this relationship was degree of nutrient specialization. It was conceivable that generalist bees were collecting nectar from one species and pollen from the other. Such generalists might therefore benefit by switching back and forth as they encountered each species in turn in the alternating arrangement. However, we found no evidence that that nutrient collection of specialists differed from that of generalists or that bees were using one *Tecoma* species for one nutrient and the other species for the other nutrient (Results, Fig. 7). This is perhaps unsurprising, given the two species used here belong to the same genus. Somme et al. (2015) reported that individual bumble bees in the field visited multiple plant species because some species were best for nectar requirements and some best for pollen requirements. This would be an interesting context in which to examine the relationship between preference and switching by individuals. More generally, the literature on dietary mixing of nutritionally complementary foods offers insight into control of the optimal ratio of foods (preference) versus the rate at which food switching occurs (Raubenheimer et al. 2009; rev. Raubenheimer and Simpson 2018).

A dependence of switching behavior on spatial distribution or nutrient mixing, as presented here, raises the possibility that the preference - switching relationship is phenotypically plastic and adjusted to external environmental conditions. We might also expect the relationship to be sensitive to a forager’s internal state. Hunger level, nutritional deficits, age, and perceived predator risk are possible factors affecting the relationship between preference and switching frequency that merit future study.

### Consequences of preference and switching for the biotic resource

The distinction between preference and switching frequency is relevant not only to the forager but also to its biotic resource, as has been argued for the evolution of floral characters important in pollinator attraction (Waser 1986; Gegear and Laverty 2001; but see Janovsky et al. 2017). Low switching frequency will cause more pollen to be exported to conspecifics and more pollen to be deposited on conspecific flower stigmas (Takimoto et al. 2022; Tenhumberg et al. 2023. As noted in the Introduction, two individuals can have the same degree of preference, yet differ markedly in switching frequency (Fig. 1b, first row).

Where so-called constancy indices (including Bateman’s Index, Jacobs Index, and Modified Jacobs Index) have been used to assess switching frequency relative to preference, pollinators show variation within and between species. Nevertheless, in a number of studies, bees and butterflies switch significantly less than expected based on overall preference (Bateman 1951; Waser 1986; Lewis 1989; Gegear and Thomson 2004; Gegear and Laverty 2001, 2005; Hopkins and Rausher 2012; Ishii and Masuda 2014). In a recent field study (Bruninga-Socolar et al. 2022), this was due primarily to flowers of the same type being closer together in space than flowers of other types. However, in some laboratory or enclosure studies that control for spacing, low switching still occurs. In light of the possible benefit of low switching frequency to plant reproductive success, plant floral traits may evolve that contribute to low switching frequency (Grant 1950; Waser 1983; Thomson et al. 2019).

Finally, the roles of preference and switching in floral evolution are not necessarily equivalent. Hopkins and Rausher (2012), studying the evolution of floral color in *Phlox drummondii*, found that preference and switching by a key pollinator, the pipevine swallowtail (*Battus philenor*), played different roles in the evolution of floral color in *Phlox* populations. The butterfly’s pattern of color preference accounted for the maintenance of floral color differences between allopatric *Phlox* populations. In contrast, the low frequency of switching by the butterfly as measured with a Bateman’s index (that is, floral color constancy) accounted for the maintenance of a color polymorphism within *Phlox* populations (Hopkins 2022).

### A closing note on floral constancy

We used the term ‘flower constancy’ (aka ‘floral constancy’) sparingly throughout this paper, despite a long history of usage in the pollination literature. We did so because the term has taken on multiple implicit or explicit meanings over the years. In earliest usage, dating back at least to the 1880’s (Bennett 1883; Christy 1883), constancy referred essentially to a strong, even absolute preference by a pollinator for one flower type in a foraging bout. Absolute preference of this kind, without using the word ‘constancy’, had been reported for bees long ago by Darwin and Müller, and even earlier by Aristotle, and was later studied by Forel and von Frisch among others (rev. Grant 1950).

Waser (1986) advocated persuasively for a focus on transitions between flower types and recommended use of Bateman’s (1951) constancy index to assess fidelity in transitions, thus putting emphasis on switching frequency. It is important to note that Bateman’s index and other constancy indices, such as the Modified Jacobs Index used here, will be correlated with common preference measures, but are not equivalent to those measures. In fact, as we have attempted to show here, the indices are particularly useful in understanding how switching frequency varies in relation to overall preference.

The term ‘flower constancy’ remains popular. Web of Science reports 248 publications from 2020-2023 referring to flower constancy or floral constancy. Recent usage is mixed. Most often, constancy is equated to preference, with preference quantified as it was quantified in our study. Sometimes constancy is assessed using Bateman’s or Jacobs indices; even then, the interpretation of index value may be made in terms of preference, not in terms of switching frequency relative to preference. The definition of constancy aside, our results suggest that ecologists should consider preference, switching frequency, and their relationship in study systems. We hope that our findings illustrate that these traits are expected to be intrinsically related but that the relationship can vary in ways important both to the ecology of the forager and that of its biotic resource.

## Acknowledgements

A discussion group at the University of Arizona encouraged us to think more about the topic of preference and switching; we thank group members Heather Briggs, Carla Essenberg, Sarah Richman, and Gordon Smith. Minjung Baek and Jacob Francis are also thanked for discussions. Robin Hopkins is gratefully acknowledged for sharing insights on variation in preference and switching.

