## Supplemental Tables 1 and 2 for "The Relationship Between Preference and Switching in Flower Foraging by Bees"

*for* Behavioral Ecology and Sociobiology

*by* Daniel R. Papaj[1] and Avery L. Russell[2]

[1] Department of Ecology and Evolutionary Biology, University of Arizona, Tucson, Arizona 85721 U.S.A.

[2] Department of Biology, Missouri State University, Springfield, MO, 65897 U.S.A.

Correspondence: DR Papaj, Department of Ecology and Evolutionary Biology, 1041 E. Lowell Street, University of Arizona, Tucson, AZ 85721, U.S.A.

**Table S1.** Species preference for each test bee. Also shown is the result of a Chi-Square Goodness-of-Fit test on each bee. Total N refers to total number of visits to flowers by the test bee. Color coding indicates species preference in absolute terms. Orange indicates statistically significant *T. alata* preference, yellow signifies significant *T. stans* preference, and white indicates no statistically significant preference at p = 0.05.*

| Bee ID | Total No. Visits | % *T. alata* Preference | Species Preferred | Chi Square Value | p value |
|---|---|---|---|---|---|
| P3 | 49 | 63 | *T. alata* | 2.94 | 0.086 |
| P5 | 50 | 26 | *T. stans* | 8.82 | 0.003 |
| P6 | 41 | 2 | *T. stans* | 35.22 | <0.0001 |
| P10 | 24 | 96 | *T. alata* | 18.38 | <0.0001 |
| P14 | 40 | 73 | *T. alata* | 7.22 | 0.007 |
| P17 | 40 | 95 | *T. alata* | 30.62 | <0.0001 |
| P19 | 21 | 38 | *T. stans* | 0.76 | 0.38 |
| P20 | 11 | 82 | *T. alata* | 3.28 | 0.07 |
| P23 | 75 | 49 | *T. stans* | 0 | 1 |
| P31 | 41 | 2 | *T. stans* | 35.22 | <0.0001 |
| P32 | 40 | 0 | *T. stans* | 38.02 | <0.0001 |
| O34 | 44 | 2 | *T. stans* | 38.2 | <0.0001 |
| O36 | 34 | 38 | *T. stans* | 1.44 | 0.23 |
| O37 | 20 | 30 | *T. stans* | 2.46 | 0.12 |
| P38 | 29 | 21 | *T. stans* | 8.82 | 0.003 |
| O39 | 9 | 56 | *T. alata* | n/a | n/a |
| O41 | 51 | 41 | *T. stans* | 1.26 | 0.26 |
| O42 | 35 | 23 | *T. stans* | 9.26 | 0.002 |
| O44 | 14 | 50 | 0 | 0 | 1 |
| O45 | 55 | 49 | *T. stans* | 0 | 1 |
| O47 | 57 | 54 | *T. alata* | 0.28 | 0.6 |
| O49 | 44 | 0 | *T. stans* | 42.02 | <0.0001 |
| O51 | 16 | 25 | *T. stans* | 3.06 | 0.08 |

* Bee O39 had too few total visits (N=9) to test with a chi-square test. A binomial test indicates that the one-tailed probability of a preference of 5 out of 9 (= 0.56) or greater, given a null expectation of 0.5 preference, is 0.5.

**Analysis of nectar collection**

Analysis of nectar collection confirmed that bees were successfully extracting nectar from both *Tecoma* species. The two *Tecoma* species differed significantly in the mean volume of nectar in untested flowers collected at the same time as test flowers, with *T. alata* having significantly more nectar than *T. stans* (Table S2; Wilcoxon Signed Rank Test, W=39, p=0.0005; N=12 sets of flowers). As expected, if bees were successfully extracting nectar, test flowers of each species consistently had significantly less nectar than time-matched control flowers (Table S2; Wilcoxon Signed Rank Tests; *T. stans*, W=36, p<0.003; *T. alata*, W=26, p<0.05; N = 12 sets of each species). Proportionately more nectar was left in the *T. alata* flowers than the *T. stans* flowers, probably because bees cannot reach far enough into the longer, narrower corollas of *T. alata* to remove the remaining nectar.

We also measured the concentration of nectar for a sample of flowers of each species, using the Brix method. The Brix method assumes all dissolved solids are sucrose, but yields good estimates of % total dissolved solids even when the solution is not 100% sucrose. Nectar of *T. stans* was more concentrated than that of *T. alata*: *T. stans*, mean concentration = 50.13% ($\pm$ 0.76 S.E.), N=15 flowers; *T. alata*, mean concentration = 33.54% ($\pm$ 0.72 S.E.), N=13 flowers. This difference was highly significant (t-test, $t_{26}$=15.67, p<0.0001).

**Table S2.** Nectar volume data for each *Tecoma* species. N = 12 sets of flowers for each species.

| **Flower Species and Category** | **Mean (± S.E.) (ul)** | **Median (ul)** | **% Flowers Empty** |
|---|---|---|---|
| ***Tecoma stans*** | | | |
| Test | 0.41 (0.30) | 0.07 | 25.0 |
| Matched Controls | 1.48 (0.26) | 0.41 | 0 |
| ***Tecoma alata*** | | | |
| Test | 5.57 (0.88) | 5.44 | 0 |
| Matched Controls | 7.58 (0.72) | 6.01 | 0 |
